# Chronic Inhaled Benzene Exposure Exacerbates Atherosclerosis in LDL Receptor-Null Mice

**DOI:** 10.1101/2025.01.29.635519

**Authors:** Igor N. Zelko, Marina V. Malovichko, Samantha A. McFall, Breandon S. Taylor, Hong Chen, Daniel J. Conklin, Sanjay Srivastava

## Abstract

**Background:** Benzene is a ubiquitous environmental pollutant generated by a variety of natural and anthropological sources. It is a known carcinogen and hematopoietic toxin; however, little is known about benzene’s potential atherogenicity.

**Hypothesis:** Inhaled benzene induces atherogenesis by increasing vascular inflammation in LDL receptor Knockout (LDLR-KO) mice.

**Methods:** Male LDLR-KO mice were exposed to HEPA-filtered air or benzene (1 ppm, 6h/day, 5days/week) for 24 weeks. For the last 12 weeks of exposure, the mice were maintained on a western diet. The single nuclei RNA sequencing (snRNAseq) of aortae was performed at Novogene. For *in vitro* experiments, splenic naïve T cells were exposed to 1 µM of hydroquinone (HQ) for 24 hours, and intracellular ROR-gamma levels were measured by flow cytometry.

**Results:** Benzene inhalation increased the aortic valve lesion area by more than 25% (P<0.05) in LDLR-KO mice. Using snRNAseq, eleven major cell types were detected, including T cells and vascular smooth muscle cells (VSMC). Benzene increased the number of T cells by 2.5-fold, proliferating T-cells by 5.8-fold, and VSMC by 1.6-fold, suggesting increased cellularity and reduced plaque stability. In addition, benzene upregulated Th17 polarization marker *Rorc* and negative regulators of apoptosis *Rag1* and *Bcl11b* while significantly attenuating the expression of proliferation inhibitor *Ms4a4b* in T cells. In VSMC, benzene downregulated extracellular matrix organization genes and upregulated platelet degranulation pathways. Polarization of T cells into Th17 was confirmed by HQ-dependent upregulation of ROR-gamma *in vitro*.

**Conclusion:** Our data suggest that inhaled benzene exposure compromises plaque cellularity and stability by facilitating T-cell proliferation and polarization, which coincides with the degradation of smooth muscle extracellular matrix and platelet activation.

---

Environmental factors, especially air pollution, which accounts for more than six million premature deaths, is associated with an increased risk for cardiovascular disease (CVD). Both human and experimental animal studies demonstrate that particulate matter (PM), especially PM_2.5_, exacerbates CVD^1^. However, less is known about the contribution of other airborne toxicants, such as volatile organic compounds (VOCs; e.g., benzene, xylene, acrolein, etc.), in air pollution-induced CVD. Benzene, a well-known human carcinogen ranked 6th on the Agency for Toxic Substances and Disease Registry (ATSDR) priority list, is abundant in petroleum products, cigarette smoke, and wildfire smoke; and is one of the top 20 chemicals generated by industrial sources in the United States^2^. Moreover, household products (e.g., detergents, paints, furniture wax, etc.) also emit considerable benzene^2^. Furthermore, people living near hazardous waste sites containing benzene and trichloroethylene are at increased risk for stroke^3^. However, little is known about the effects of benzene exposure on atherosclerosis – a major contributor to myocardial ischemia and stroke. Thus, in this study, we examined the effect of chronic benzene inhalation on atherosclerosis in LDL receptor-null mice.

LDLR-null mice were exposed to 1 ppm benzene (reflecting moderate human exposure) or HEPA-filtered air for 24 weeks, as described in Fig. 1, and atherosclerotic lesions were quantified in the aortic valves. Chronic benzene exposure significantly increased atherosclerotic plaque formation (**Fig. 1A**) without affecting total plasma cholesterol (data not shown). Importantly, benzene (1 ppm) exacerbated atherogenesis at levels 50-fold lower than in tobacco smoke or in occupational exposures and 300-fold lower than that required for hematopoietic toxicity in experimental animals. Lesion composition and vascular inflammation of the aortae of benzene-exposed mice were examined by single-nuclei RNA sequencing (snRNAseq) of the aortae (after carefully removing the adventitia) using the 10x Genomics Gemcode Technology at Novogene Inc. The snRNAseq generated 3.1×10^9^ reads for air-exposed, and 3.4×10^9^ reads for benzene-exposed groups. There were 11 major cell types: endothelial cells (ECs; *Cdh5, Pecam1*), vascular smooth muscle cells (VSMCs; *Myh11, Mylk*), monocytes and macrophages (Mo/Mf; *Lgals3, Adgre*), T-cells (*Cd4, Cd8a, Themis*), fibroblasts (Fibro; *C7, Abca8a*), B-cells (*Cd79a, Cd79b*), Schwann cells, neurons, and mesothelial cells (Schwann/Neuro; *So×10, Cdh19, Cadm2, Upk3b*), adipocytes (Adipo; *Pck1, Adipoq*), erythroid (*Hba-a1, Hba-a2*), and proliferating cells (Proliferating; *Top2a, Mki67, Diaph3*). Aortic cells were clustered into 23 sub-populations in both air- and benzene-exposed mice. Benzene exposure robustly increased the abundance of lesional T-cells, VSMC, and Proliferating cells (**Fig. 1B**). Re-clustering of proliferative cells revealed that benzene exposure increased the abundance of lesional T-cells (*Cd4, Cd8a, Themis*; inset: **Fig. 1B**). Benzene exposure facilitated the shift of T-cells toward pro-atherogenic CD8^+^ and follicular helper (Tfh) phenotype positive in *Bcl6* expression (**Fig. 1Ci**). Reactome pathway analysis supported that benzene exposure robustly affected T-cell receptor (TCR) signaling (**Fig. 1Cii**). Benzene shifted T-cell regulatory phenotype towards Th17 as evidenced by marked induction of the RAR-related orphan receptor gamma (*Rorc*) gene (**Fig. 1Di**). *In vitro*, incubation of murine spleen-derived CD4^+^ thymocytes with the benzene metabolite hydroquinone (1 µM) upregulated ROR-γ protein as assessed by flow cytometry (**Fig. 1Dii**). To examine the potential cell-cell communications (based on ligand-receptor interaction) in the lesion, we performed CellChat analysis. Benzene exposure appreciably increased the ligand-receptor interaction strength while slightly reducing the number of inferred interactions (**Fig. 1Ei and 1Eii**). Additionally, benzene exposure significantly changed information flow for at least 35 of the 229 analyzed pathways (**Fig. 1Eiii**). Importantly, benzene exposure induced marked perturbations in IL-16 (a chemoattractant and ligand for CD4) signaling between lesional T-cells and other cell types, e.g. macrophages, VSMC, and fibroblasts (**Fig. 1Fi**). Benzene exposure also increased communication probabilities between lesional lymphatic endothelial cells (EC_5 – Lyve1, Mmrn1, and Prox1) and T-cells (**Fig. 1Fii**).

**Figure 1:**
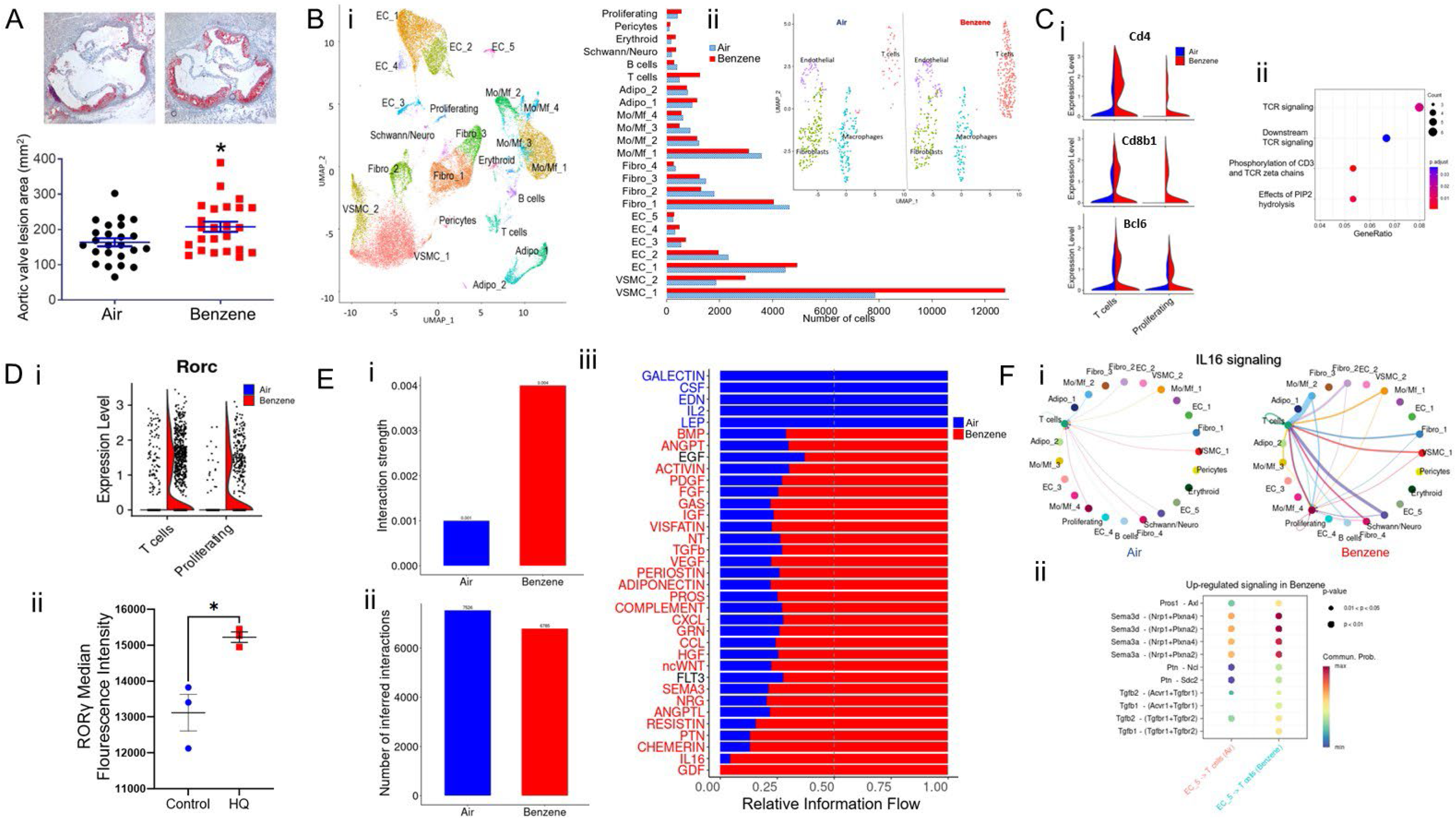
Atherogenic effects of benzene. Eight-week-old male LDL receptor-null mice maintained on normal chow were exposed to HEPA-filtered air (Air) or benzene (1ppm, 6h/d, 5d/week) for 12 weeks. The mice were then fed a Western diet for another 12 weeks of exposure. **A**. *Lesions in the aortic valves*. Lesional lipids were stained with Oil Red O. Values are mean ± SEM, *p<0.05. **B**. *Lesion composition* - i) clustering of cells in the aortae of air-exposed mice as assessed by snRNAseq, and ii) relative distribution of cells in the aortae of air (blue, n=6) and benzene (red, n=6) exposed mice. Inset – reclustering of proliferative cells in the aortae. EC - Endothelial cells, VSMC - Vascular smooth muscle cells, Fibro - fibroblasts, Mo/Mf - Monocytes/macrophages, Adipo – Adipocytes. snRNAseq analysis was performed with Seurat. **C**. *T-cell-specific gene expression* – i) upregulated expression of Cd4, Cd8, and Tfh marker Bcl6; ii) Reactome pathway analysis of up-regulated genes. **D**. *RAR-related orphan receptor -* i) RAR-related orphan receptor transcription of lesional T-cells and proliferative cells in benzene-exposed mice and ii) flow cytometric analysis of RAR-related orphan receptor gamma (RORγ) protein in murine spleen-derived CD4^+^ T-cells incubated with benzene metabolite, hydroquinone (HQ, 1µM, 18h). Values are mean ± SEM, *p<0.05. **E**. *Cell-chat analysis -* i) the mean interaction between cells, (ii) the number of inferred interactions, (iii) relative information flow of significant signaling pathways. **F**. *Effect of benzene on select signaling pathways -* i) IL-16 signaling network and ii) upregulated signaling between lymphatic EC and T-cells.

This is the first study to report the exacerbation of atherosclerosis by benzene exposure. The snRNAseq data clearly demonstrates that benzene exposure affects lesion inflammation likely via altered T-cell abundance and signaling. T-cells are present in all stages of plaque development and are critical modulators of atherogenesis. Consistent with previous reports on single-cell RNAseq in murine atherosclerotic plaques^4^, we observed clusters of CD8^+^, CD4^+^ Th17, Tfh, and CD4^+^CD8^+^ cells, whereas markers of T_reg_ and Th1 were difficult to detect in the aortae of both the air- and benzene-exposed mice. Increased transcription of *Rorc* in the aortae of benzene-exposed mice reflects the induction of mostly pro-atherogenic Th17 subtype while upregulating expression of Bcl6 indicates a transition of T-cells towards pro-atherogenic follicular helper (Tfh) cells^5^. Benzene exposure also increased the number of CD8^+^ cells that may promote necrotic core formation and destabilize plaques. Induction of IL-16 signaling may augment T-cell recruitment in the plaques of benzene-exposed mice. Moreover, enhanced expression of TGFβ in lymphatic endothelial cells may promote Th17 differentiation in benzene-exposed mice. Additional studies are required to rigorously examine how benzene and its biotransformation affects atherogenesis and vascular inflammation and whether these effects are dose and sex dependent.

## Acknowledgement

The authors acknowledge the technical assistance by Ms. Millicent Winner and Ms. Fangping Yuan. This research was supported in part by NIH grants ES023716, R01 ES033531, R01 HL149351 (to SS), R01 HL156362 and R01 HL146134 (to SS and HC), R01 HL171763 (to DJC), and R21ES033334 (to INZ).

## Notes

Conflict of interest statement: None of the authors have any conflict of interest.

### Competing Interest Statement

The authors have declared no competing interest.

